# Multidrug resistance and high prevalence of class 1 integrons in *Escherichia coli* isolated from waters and vegetables in Nsukka and Enugu, Nigeria

**DOI:** 10.1101/2020.01.06.895748

**Authors:** Chinyere B. Chigor, Ini-Abasi I. Ibangha, Nkechinyere O. Nweze, Chizoba A. Ozochi, Valentino C. Onuora, Yinka Titilawo, Tatyana N. Chernikova, Peter N. Golyshin, Vincent N. Chigor

**Affiliations:** Water and Public Health Research Group (WPHRG), University of Nigeria, Nsukka, Enugu State, Nigeria; Department of Plant Science and Biotechnology, Faculty of Biological Sciences, University Nigeria, Nsukka, Enugu State, Nigeria; Department of Microbiology, Faculty of Biological Sciences, University of Nigeria, Nsukka, Enugu State, Nigeria; Department of Biology/Microbiology/Biotechnology, Alex Ekwueme Federal University, Ndufu-Alike Ikwo, Ebonyi State, Nigeria; School of Natural Sciences, Bangor University, Bangor Gwynedd, United Kingdom

**Author notes:** Corresponding author (VNC). “VNC and PNG are Joint Senior Authors”.

**Keywords:** multidrug resistance, Integrons, pathogenic *E. coli*, wastewater-irrigated vegetables, public health risks

## Abstract

In spite of treated wastewater presenting itself as an attractive alternative to scarce quality water in the developing countries, the associated contamination of fresh produce by irrigation waters leading to outbreak of foodborne illnesses is on the rise. Horizontal transfer of integrons play important role in the spread and maintenance of antimicrobial resistance among strains of *Escherichia coli*. This study assessed the effluents from the University of Nigeria, Nsukka Wastewater Treatment Plant (UNN-WWTP) as well as vegetables irrigated with the effluent, and vegetables sold in selected markets from Nsukka and Enugu cities for the presence of *E. coli* and determined the prevalence integrons in multidrug-resistant isolates. Isolation of *E. coli* was done using eosin methylene blue agar and isolates subjected to Gram staining for identification of presumptive colonies. Confirmation of *E. coli* was achieved by polymerase chain reaction (PCR) technique, targeting beta-glucuronidase (*uid*A). Resistance to antibiotics was determined using the Bauer-Kirby disk diffusion assay and the Clinical and Laboratory Standard Institute criteria. Integrons were detected by multiplex PCR using primers specific for class 1 and 2 integrons. A total of 178 *E. coli* isolates were obtained from WWTP effluent (41), and vegetables from greenhouse (46), farms (55) and market (36). Multi-drug resistance was detected in all the isolates, ranging from five-drug resistance in a single isolate to 16-drug resistance patterns in two different isolates. Of the total isolates, class 1 integrons were abundantly detected in 175 (98.3%) and class 2 in 5 (2.8%). All the class 2 integrons were found in isolates that were positive for class 1. The high detection of *E. coli* in the studied effluent and vegetables pose potential public health hazards heightened by observed multidrug resistance in all the isolates and the high prevalence of class 1 integron. It is concluded that the vegetable samples are significant reservoirs for potentially pathogenic *E. coli.* Therefore, vegetable irrigation farming with unsafe water should be discontinued, while appropriate improvement strategies to ensure compliance should be facilitated without further delay.

## Introduction

Pathogenic *Escherichia coli* causes significant morbidity and mortality worldwide [1–3]. Reported risk factors in the developing countries and sub-Saharan African regions include poor hygiene, unsafe water, improper disposal of waste and faeces, and contaminated food, local beverages and vegetables [2, 4, 5]. Vegetables can become contaminated with pathogenic and commensal bacteria from animals and humans, during growth, harvesting, distribution, storage and processing [6]. Although the contamination of fresh produce by irrigation waters has led to outbreak of foodborne illnesses, yet treated wastewater presents itself as an attractive alternative to scarce quality water in the developing countries.

*E. coli* has been reported as an aetiological agent of diarrhoea in both the northern and south-western parts of Nigeria [7–12]. In a study that detected *E. coli* in 119 (44.74%) of 270 diarrhoeal stool samples in Enugu and Onitsha cities, south-eastern Nigeria [13], enterotoxigenic *E. coli* (ETEC) was reported as the second most prevalent pathotype (21.57%) after enteropathogenic *E. coli* (EPEC) (49.02%). Likewise, a study conducted in Nsukka, that involved watery stools, drinking water, and some fruits and vegetables collected during the rainy periods (between April and October) over 3 year sampling regime (1996 to 1998), [4] reported that enteropathogenic *E. coli* (EPEC) was detected in 9 (1.8%) of 500 stool samples, whereas no enteric bacterial pathogen was isolated from the fruits and vegetables. There appears to be no reports on ETEC prevalence in humans and on irrigated vegetables in Nsukka.

Excessive and inappropriate usage of antimicrobials in preventing or treating human and veterinary bacterial infectious diseases has led to increased antimicrobial and multidrug resistance (MDR) and the risk of transmission of antibiotic resistant bacteria (ARB) and antibiotic resistant genes (ARGs) from one country to another is a growing global challenge. [14–16] Attention should be given to how anthropogenic activities might be causing evolution of antibiotic resistance in the environment [16], and studies have shown that waste water treatment plants form a significant reservoir of resistance genes and suggested that waste water disposal increases the reservoir of resistance determinants in the environment either by the addition of resistance genes or input of agents selective for resistant phenotypes [17].

Along with transposons and plasmids, integrons, genetic elements commonly found in bacterial genomes that allow efficient acquisition and expression of exogenous genes, are central in the dissemination of antibiotic resistance among Gram-negative bacteria [18,19]. Horizontal transfer of integrons have been shown to play important role in the spread and maintenance of antimicrobial resistance among strains of *E. coli* and ARB can be transferred across borders by human travelers, animal and insect vectors, agricultural products and surface water [15,20]. Not much is known about the risk factors in spreading across local borders.

It is thought that University towns, characterized by regular and significant demographic changes arising from admissions and vacations, could play major role in dissemination of resistance determinants locally, and even internationally where the institution has a good number of internationals. Nsukka, in Southeast Nigeria, is the location of one of Nigeria’s biggest universities and one also in which the town developed around the university. This study assessed the effluent from the University of Nigeria, Nsukka Wastewater Treatment Plant (UNN-WWTP) as well as vegetables irrigated with the effluent and vegetables sold in selected markets for the presence of *E. coli* and determined the prevalence integrons in multidrug-resistant isolates.

## Methods

### Description of study area

The university town of Nsukka (6.8429° N, 7.3733° E) is in Enugu State, southeast Nigeria, with an area of 1,810 km^2^ and a population of 309,633 (NPC 2006). The sewage treatment facility (WWTP) in Nsukka, consisting of a screen, primary settling (Imhoff) tank, sludge drying beds and two oxidation ponds, is situated at the northwest end of the University of Nigeria, Nsukka. The final effluents have been widely utilized for fresh produce irrigation during dry season.

### Cultivation of *Amaranthus* in the greenhouse

The most commonly cultivated vegetables in the study area, during the dry season, include the green leafy vegetable amaranth (*Amaranthus* spp), fluted pumpkin leaves (*Telfaria occidentalis*), scarlet eggplant leaf (*Solanum aethiopicum*) and water leaf (*Talinum fruticosum*). In this study, *Amaranthus* was chosen, being the second most produced and sold leafy vegetable, after *Telfaria* [21], and it eqaully grows very easily and matures faster. Amaranths were grown for 10 weeks (July 26 to October 03, 2014) in earthen pots at the Soil Science Departmental greenhouse. They were irrigated daily using the sprinkler method. A total of 60 earthen pots were used for the cultivation of vegetables, 48 were irrigated with treated wastewater (final effluent of the University of Nigeria, wastewater treatment plant (WWTP) and 12 with tap water. The pots irrigated with tap water served as the control.

### Collection of Samples

Samples collected for this study included treated wastewater and vegetables. Sampling was done according to the standard procedure [22]. Effluents were collected with 10 L plastic cans for irrigation of the green house vegetables. Samples of the WWTP effluent were collected using sterile wide-mouthed, screw-capped 250-ml bottles. Vegetables were obtained from the green house, irrigated gardens and local markets in Nsukka and Enugu metropolis, during December 2014. Samples of the major vegetables cultivated during the dry season include fluted pumpkin leaves (*Telfaria occidentalis*), scarlet eggplant leaf (*Solanum aethiopicum*), water leaf (*Talinum fruticosum*) and the green vegetable (*Amaranthus* Spp) were collected. All samples were transported on ice to the laboratory and analysed within 6 h of collection.

### Isolation and identification of presumptive *E. coli*

This was carried out at the Water and Public Health Laboratory, University of Nigeria Nsukka. Exactly 5 g of each vegetable sample was homogenized in a clean porcelain mortar, and 1 g of the homogenate diluted into 9ml normal saline [23, 24]. Serial dilutions (10-fold) were made by pipetting out 1ml stock solution into successive 9ml of sterile normal saline bottles. A 1 ml working sample dilution (10^−1^and 10^−2^) was spread-plated onto eosin methylene blue (EMB) agar (Oxoid, UK), incubated at 44 °C for 18-24 h. Raised, entire colonies with dark greenish metallic sheen, typical *E. coli* colonies were subjected to Gram-staining [25] and standard biochemical tests (IMViC). All presumptive *E. coli* isolates were sub-cultured in tryptic soy broth (Oxoid, UK) and then stored at −20 °C for further investigations. All media were prepared following the manufacturers’ instructions.

### Extraction of genomic DNA

Genomic DNA were extracted from a pure culture of each isolate grown overnight on nutrient agar at 37°C, by the conventional boiling method, as described [28]. Briefly, one loopful of bacterial cells was suspended in 1ml of sterile distilled water. The bacterial suspensions were then heated for 5 min at 100°C, cooled to room temperature and centrifuged at 12,000 xg for 5 min to remove the debris. The supernatant was stored at −20°C and used as the template DNA for PCR analysis.

### Detection of beta-glucuronidase (*uid*A) gene for confirmation of *E. coli*

The confirmation of *E. coli* was achieved by polymerase chain reaction (PCR) detection of the target beta-glucuronidase (*uid*A). This was done at the School of Natural Sciences, Bangor University, United Kingdom, following the procedures described [29, 30]. The extracted DNA were cleaned using QIAGEN (QIAEX®II) gel extraction kits and kept at −20°C. PCRs were carried out with BIORAD DNA Engine Tetrad®Peltier Thermal Cycler (BIORAD, USA). The PCR reaction mixtures consisted of 25 μl of PCR Master Mix (Thermo Scientific, (EU) Lithuania), 0.5 μl each of oligonucleotide primers (Eurofins Genomics, Ebersberg Germany), 10 μl of template DNA and 14 μl of nuclease free water to constitute a total reaction volume of 50 μl. The PCR cycling conditions, with some modifications, were in accordance with the protocols prescribed elsewhere [31]. *E. coli* strain (NCTC 13353) and *Enterobacter aerogenes* (NCTC 10006) were used as positive and negative controls respectively for *E. coli* genus identification. The oligonucleotide sequence of primers used, target genes and expected amplification products are given in Table 1. For gel electrophoresis, 3 μl of DNA ladder (1 Kb Plus DNA Ladder; Invitrogen), 6 μl of positive control and 10μl of template DNA were ran on 2.5% (w/v) agarose gels in 1x-TBE buffer (0.09 M Tris-borate and 0.002 M EDTA, pH 8.0) at 100V for 25-30 min. The gels were viewed and photographed with BIORAD Molecular Imager® Gel Doc™ XR Imaging System (BIORAD, USA).

**Table 1:**
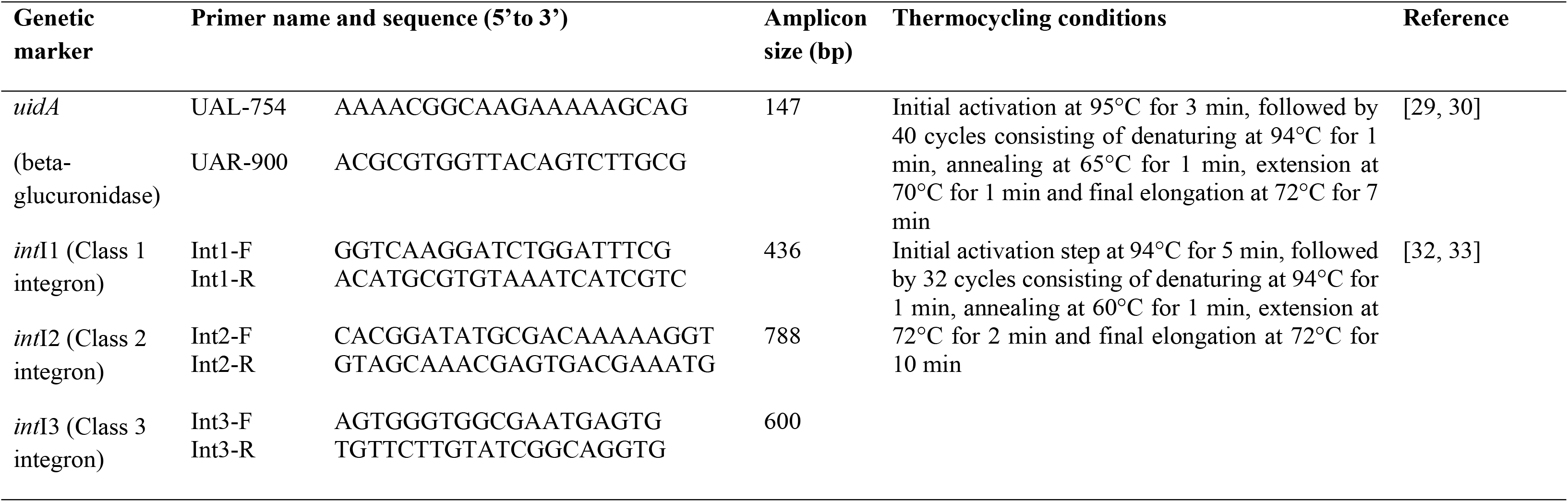
Primers for the detection of *E. coli* and integrons.

### Antibiotic susceptibility testing

Isolates were subjected to antibiotic susceptibility testing using the Kirby-Bauer disc diffusion test [26]. Evaluation of results was based on the standards of the Clinical Laboratory Standards Institute (CLSI) [27]. Briefly, isolates grown on nutrient broth were suspended into sterile normal saline (0.9% (w/v) NaCl) with the aid of a sterile wire loop until the turbidity equivalent of 0.5 McFarland standard was reached. Sterile non-toxic cotton swabs were dipped into the standardized inoculum and used to smear the entire surface of the Muller-Hinton agar (Thermo Fisher Scientific, USA) plates. Antibiotic discs were placed aseptically using sterile forceps. All plates were incubated at 35±2oC for 16 to 18 h. The following antibiotics were employed for the test: Amoxycillin (AMX) 10μg, Ampicillin (AMP) 10μg, Metronidazole (MTZ) 5μg, Rifampicin (RIF) 5μg, Vancomycin (VAN) 30μg, Cloxacillin (COX) 5μg, Penicillin G (PNG) 10iu, Streptomycin (STR) 10μg, Erythromycin (ERT) 15μg, Clarithromycin (CLR) 15μg, Cefuroxime (CXM) 30μg, Chloramphenicol (CHL) 30μg, Imipenem (IPM) 10μg, Tetracycline (TET) 30μg, Ciprofloxacin (CIP) 5μg, Trimethoprim (TMP) 5μg, Norfloxacin (NOR) 10μg, Sulphamethoxazole (SMZ) 25μg. The *E. coli* ATCC 25922 strains was used as control for antibiotic susceptibility testing. Zones showing complete inhibition around the discs were measured and classified as resistant (R), intermediate (I) and susceptible (S) according to the diameters of the zones recorded to the nearest millimetres.

### Detection of integrons

The isolates were screened for class 1, 2 and 3 integrons by a multiplex PCR procedure as described by Machado *et al.* [32] and Karger *et al.* [33]. The PCR reactions (a total volume of 50μl reaction mixture) each consisted of consisting of 10 μl Buffer of 5x MyTaq Reaction Buffer (Bioline, with dye), 0.75μl of each the primers *int*I1, *int*I2 and *int*I3 (Eurofins Genomics, Ebersberg, Germany).), 27.25μl nuclease free water (Sigma-Aldrich), 0.25μl MyTaq DNA polymerase (Bioline) and 5μl DNA template. For gel electrophoresis, 3μl of DNA ladder (1 Kb Plus DNA Ladder; Invitrogen), 6 μl of positive control and 10 μl of samples were ran on 1.5% (w/v) agarose gels in 1x-TBE buffer (0.09 M Tris-borate and 0.002 M EDTA, pH 8.0) at 100 V for 25 min. The gels were viewed and photographed with BIORAD Molecular Imager® Gel Doc™ XR Imaging System.

## Results and Discussion

It is worthy of note the amaranths irrigated with wastewater effluent were of higher yields compared to the controls irrigated with tap water. This is attributable to the fact the effluent is rich in nutrients. It also underscores the continued preference of effluents over the scarce treated water by the vegetable farmers. Despite a perceived understanding farmer have on the use of unsafe WWTP effluent, this knowledge seems not bother them.

In the present study, the total number of samples collected were 288 including WWTP effluents (60), greenhouse (60), farm (84) and market (84) vegetables. A total of 178 *E. coli* isolates were confirmed by PCR amplification of the β-glucuronidase *uid*A gene (Fig 1), with 41 isolates from the WWTP effluents, 46 greenhouse from the effluent irrigated vegetables, 55 from vegetables collected from gardens that produce vegetables sold in local markets and 36 from vegetables bought from selected markets in Nsukka and Enugu (Table 2).

**Fig 1.**
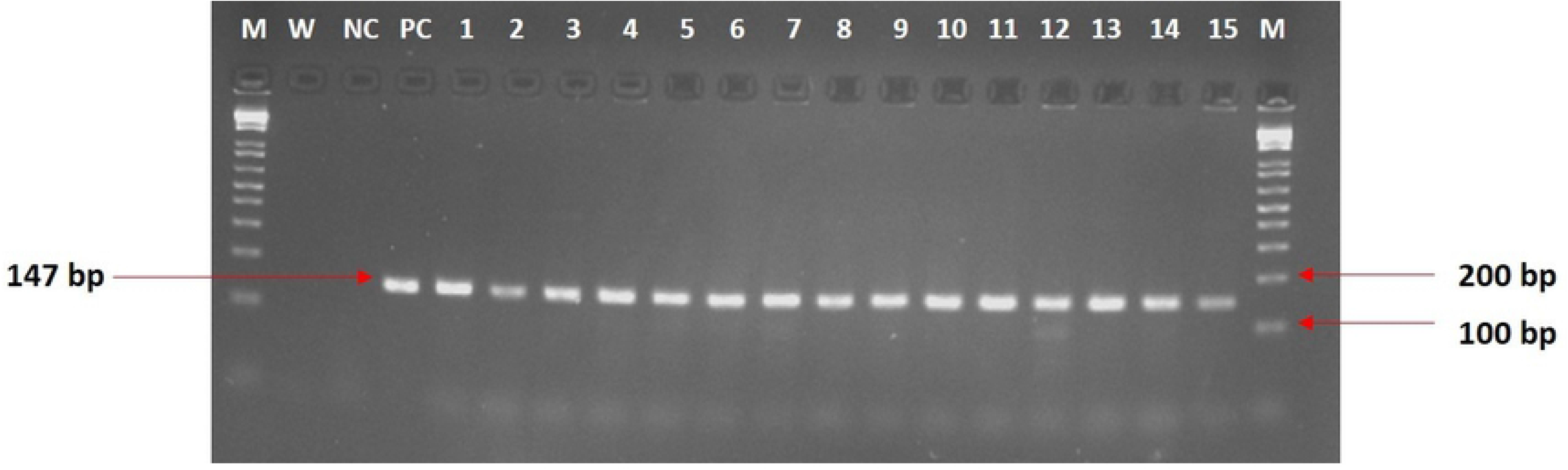
PCR products for *E. coli* confirmation by *uid*A gene amplification. This is the Fig 1 legend: M: Molecular weight marker (1 KB), W: water, NC: Negative control (*Enterobacter aerogenes*), PC: Positive control (*E. coli*; NCTC 13353), Lanes 1-15: Positive isolates

**Table 2:** Isolation of *E. coli* in effluent wastewater and vegetable samples.

| Sample type | No. of samples | No. of *E. coli* isolates |
| --- | --- | --- |
| WWTP effluent | 60 | 41 |
| Greenhouse vegetables | 60 | 46 |
| Farm vegetables | 84 | 55 |
| Market Vegetables | 84 | 36 |
| **Total** | **288** | **178** |

Table 3 shows the antibiotics susceptibility profiles of the 178 *E. coli* isolates tested with 18 different antibiotics. Generally, higher resistance percentages were observed in *E. coli* from market vegetables compared with others. This could be attributable to further contamination of vegetables by clinical *E. coli* strains arising directly from handling by sellers. All the *E. coli* isolates, showed susceptibility to imipenem and only 5.6% (10/178) of all the isolates were resistant to cefuroxime (a cephalosporin). Chloramphenicol, ciprofloxacin and norfloxacin were very effective. The most significant resistance phenotypes were detected among sulphamethoxazole (58.4%), amoxicillin (52.8%), tetracycline (47.2%), trimethoprim (44.9%) and streptomycin (37.1%), as these antibiotics are commonly used in the studied communities (Table 2).

**Table 3:** Antibiotics susceptibility profile of *E. coli* isolates.

| S/N | Antimicrobial Agent<br>(disc concentration) | Class (Subclasses) | Code | Number of isolates resistant (%) |  |  |  |  |
| --- | --- | --- | --- | --- | --- | --- | --- | --- |
|  |  |  |  | Effluent<br>(n = 41) | Greenhouse<br>(n = 46) | Farm<br>(n = 55) | Market<br>(n = 36) | TOTAL<br>(n = 178) |
| 1 | Cloxacillin (5µg) | β-Lactam (Penicillins) | COX | 41 (100) | 46 (100) | 55 (100) | 36 (100) | 178 (100) |
| 2 | Metronidazole (50µg) | Nitroimidazoles | MTZ | 41 (100) | 46 (100) | 55 (100) | 36 (100) | 178 (100) |
| 3 | Vancomycin (30µg) | Glycopeptides | VAN | 41 (100) | 46 (100) | 53 (96.4) | 36 (100) | 176 (98.9) |
| 4 | Rifampicin (5µg) | Rifamycins | RIF | 41 (100) | 45 (97.8) | 52 (94.6) | 34 (94.4) | 172 (96.6) |
| 5 | Penicillin (10U) | β-Lactam (Penicillins) | PNG | 39 (95.1) | 45 (97.8) | 54 (98.2) | 34 (94.4) | 172 (96.6) |
| 6 | Erythromycin (15µg) | Macrolides | ERT | 40 (97.6) | 46 (100) | 50 (90.9) | 32 (88.9) | 168 (94.4) |
| 7 | Clarithromycin (15µg) | Macrolides | CLR | 40 (97.6) | 37 (80.4) | 47 (85.5) | 36 (100) | 160 (89.9) |
| 8 | Ampicillin (5µg) | β-Lactam (Cephalosporins) | AMP | 34 (82.9) | 38 (82.6) | 50 (90.9) | 36 (100) | 158 (88.7) |
| 9 | Sulphamethoxazole (25µg) | Sulphonamides | SMZ | 23 (56.1) | 27 (58.7) | 26 (47.3) | 28 (77.8) | 104 (58.4) |
| 10 | Amoxicillin (10µg) | β-Lactam (Penicillins) | AMX | 23 (56.1) | 24 (52.2) | 26 (47.3) | 21 (58.3) | 94 (52.8) |
| 11 | Tetracycline (30µg) | Tetracyclines | TET | 25 (61.0) | 20 (43.5) | 16 (29.1) | 23 (63.9) | 84 (47.2) |
| 12 | Trimethoprim (5µg) | Dihydrofolate Reductase<br>(DHFR) inhibitors | TMP | 27 (65.9) | 20 (43.5) | 12 (21.8) | 21 (58.3) | 80 (44.9) |
| 13 | Streptomycin (10µg) | Aminoglycosides | STR | 15 (36.6) | 21 (45.7) | 16 (29.1) | 14 (38.9) | 66 (37.1) |
| 14 | Chloramphenicol (30µg) | Phenicol | CHL | 4 (9.8) | 6 (13.0) | 6 (10.9) | 9 (25.0) | 25 (14.0) |
| 15 | Ciprofloxacin (5µg) | Fluoroquinolones | CIP | 5 (12.2) | 7 (15.2) | 6 (10.9) | 7 (19.4) | 25 (14.0) |
| 16 | Norfloxacin (10µg) | Fluoroquinolones | NOR | 2 (4.9) | 7 (15.2) | 9 (16.4) | 5 (13.9) | 23 (12.9) |
| 17 | Cefuroxime (30µg) | β-Lactam (Cephalosporins) | CXM | 2 (4.9) | 2 (4.4) | 2 (3.6) | 4 (11.1) | 10 (5.6) |
| 18 | Imipenem (10µg) | β-Lactam (Carbapenems) | IMP | 0 (0.0) | 0 (0.0) | 0 (0.0) | 0 (0.0) | 0 (0.0) |

Multidrug resistance (MDR) has been frequently reported in Nigeria among *E. coli* isolates obtained from human specimens [12, 37, 42, 43], animal sources [36] and environmental samples [38, 42, 44, 45]. In the present study, all the isolates were multidrug resistant, ranging from 5-drug to 16-drug resistance patterns. Although some studies have reported a high removal efficiency for total ARGs in wastewater [46], our data suggest that sewage treatment process at UNN is not effective in reducing ARGs as all the *E coli* isolated from the effluent were MDR. The spread of AMR often limits the availability of therapeutic options to only a very few efficacious antibiotics [47]. The last-resort drugs, the carbapenems such as imipenem (used in this study) and meropenem, are themselves not only increasingly challenged by emerging resistance, as evident from the data presented here, but are not affordable in the developing regions.

Multidrug resistance (MDR) was detected in all *E. coli* isolates, and although this study did not determine the full virulence potentials of all the isolates subjected to antimicrobial susceptibility testing (AST), irrigational use of WWTP effluent represents a pathway for human exposure to antibiotic-resistant commensal and pathogenic bacteria. Vegetable farming at the site should therefore be discontinued as it presents significant threat to the health of consumers of such vegetables.

It is known that AMR and MDR in *E. coli* are acquired by the transfer of mobile genetic elements, such as plasmids, transposons and integrons [15, 18, 20]. In the present study, of the 178 *E. coli* isolates, class 1 integrons were detected (Fig 2) in 175 (98.3%), and class 2 in 5 (2.8%). All the class 2 integrons were found in isolates that were positive for class 1. Such co-carriage has been previously published on *E. coli* from meat turkeys in Italy, [34].

**Fig 2.**
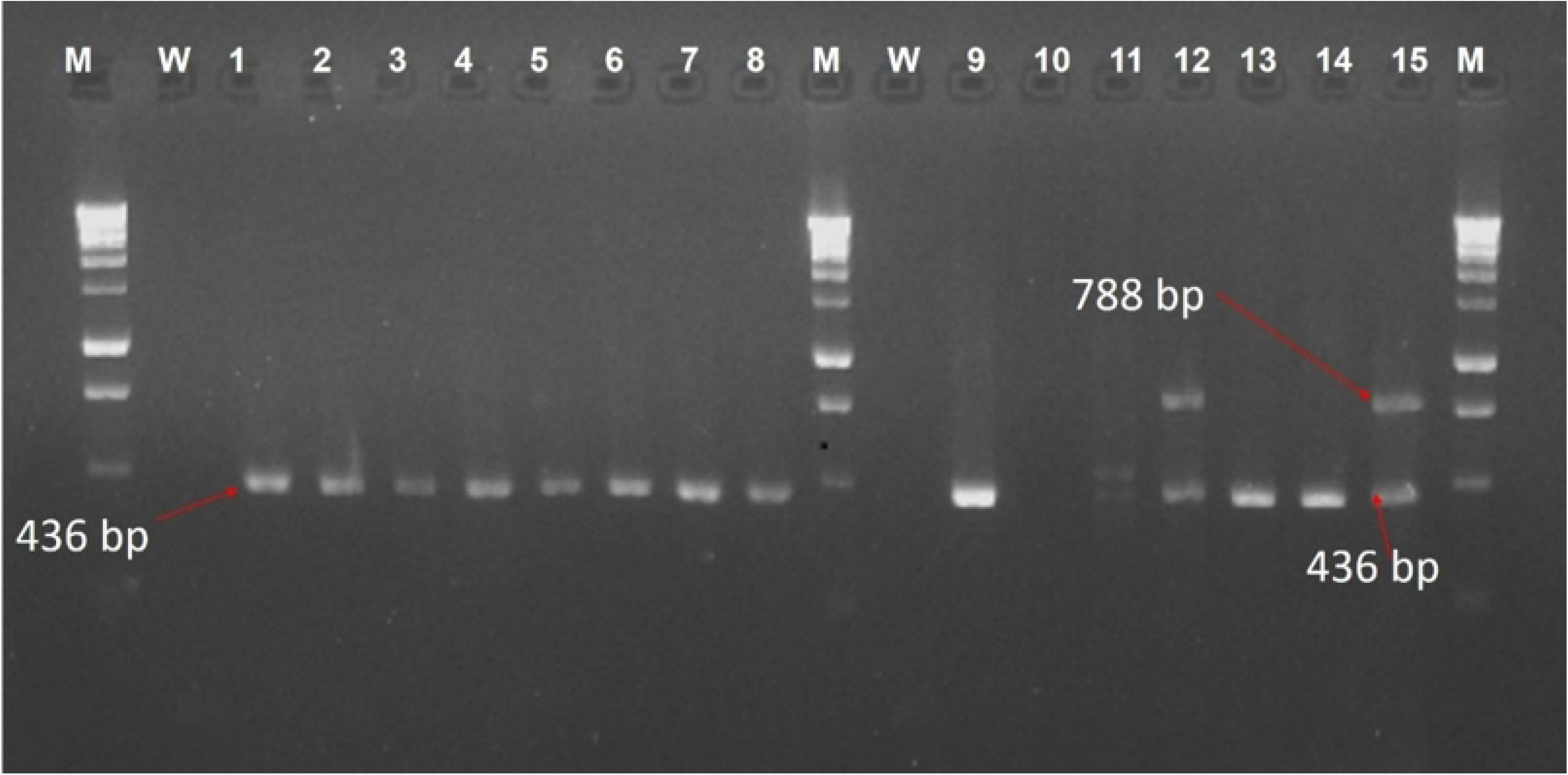
Multiplex PCR products for detection of class 1 and 2 integrons. This is the Fig 2 legend: M: Molecular weight marker (1 KB Plus ladder), W: water, Lanes 1-15: *E. coli* isolates

The integron carriage rate for the 137 vegetable isolates was 97.8%, whereas the rate for 41 effluent isolates was 100%. Considering that all the isolates were MDR, the detected high prevalence of class 1 integron is not surprising and compares with a previous study that reported that MDR phenotypes were observed in 96.8% of the integron-positive isolates [35]. These rates portend serious public health risks as it is known that class 1 integron could carry diverse antibiotic resistance genes (ARGs) and conduct horizontal gene transfer among microorganisms [20].

The data presented here shows that class 2 integrons were less frequently detected 5 (2.8%). Similar data have been published for Enterobacteriaceae in Nigeria [36, 38] and elsewhere [34, 39, 40]. Ramírez et al [39], reported that unlike the widespread distribution of class 1 integron within Gram-negative bacilli, only *Acinetobacter baumannii* and Enterobacter cloacae harboured class 2 integrons at a high frequency. However, in an earlier study in China [41], Class 2 integrons were present in 25 (80.6%) of the *Shigella sonnei* isolates and 29 (87.9%) of the *S. flexneri* isolates whereas class 1 integrons were found in only 6 (9.4%) of *Shigella* spp. isolates.

## Conclusions

The present study revealed high detection of *E. coli* in the studied effluent and vegetable samples and represent potential public health hazards intensified by observed multidrug resistance in all the isolates and the high occurrence of class 1 integrons. It is concluded that UNN-WWTP is a significant reservoir for diarrheagenic *E. coli*. Vegetable farming at the site should therefore be discouraged as it presents significant threat to the health of consumers of such vegetables.

## Acknowledgments

The authors are also grateful to Mr. Uchenna Oyeagu of the Water and Public Health Laboratory, University of Nigeria, Nsukka, Nigeria, for laboratory assistance.

